# A red herring in zebrafish genetics: allele-specific gene expression can underlie altered transcript abundance in zebrafish mutants

**DOI:** 10.1101/2021.08.06.455380

**Authors:** Richard J White, Eirinn Mackay, Stephen W Wilson, Elisabeth M Busch-Nentwich

## Abstract

In model organisms, RNA sequencing is frequently used to assess the effect of genetic mutations on cellular and developmental processes. Typically, animals heterozygous for a mutation are crossed to produce offspring with different genotypes. Resultant embryos are grouped by genotype to compare homozygous mutant embryos to heterozygous and wild-type siblings. Genes that are differentially expressed between the groups are assumed to reveal insights into the pathways affected by the mutation.

Here we show that in zebrafish, differentially expressed genes are often overrepresented on the same chromosome as the mutation due to different levels of expression of alleles from different genetic backgrounds. Using an incross of haplotype-resolved wild-type fish, we found evidence of widespread allele-specific expression, which appears as differential expression when comparing embryos homozygous for a region of the genome to their siblings. When analysing mutant transcriptomes, this means that differentially expressed genes on the same chromosome as a mutation of interest may not be caused by that mutation.

Typically, the genomic location of a differentially expressed gene is not considered when interpreting its importance with respect to the phenotype. This could lead to pathways being erroneously implicated or overlooked due to the noise of spurious differentially expressed genes on the same chromosome as the mutation. These observations have implications for the interpretation of RNA-seq experiments involving outbred animals and non-inbred model organisms.

## Introduction

Large scale genetic screens to identify gene function by randomly introducing mutations have been a staple of zebrafish genetics for several decades (Driever et al., 1996; Haffter et al., 1996; Kettleborough et al., 2013). The advent of RNA-sequencing (RNA-seq) has enabled investigators to estimate the location of such mutations in the genome, while also providing information regarding gene expression levels and affected cellular pathways in the mutants. The bioinformatics pipelines which process RNA-seq data to generate gene expression information focus on transcript abundance, differential splicing, and gene set enrichments, and, in general, genomic location is not considered when assessing genes that are differentially expressed in a mutant context. Here, we report that physical location can impact a gene’s likelihood of being differentially expressed in mutant zebrafish.

In the typical protocol for introducing new mutations, male zebrafish from a laboratory wild-type strain are treated with *N*-ethyl-*N*-nitrosourea (ENU) to mutagenise sperm (Kettleborough et al., 2011; Mullins et al., 1994). The mutagenised fish (G0) are mated with wild-type females to produce F1 offspring, each heterozygous at random novel mutation sites. F1 fish are outcrossed with wild types to produce clutches of F2 offspring, which are subsequently incrossed to produce F3 embryos. The F3 clutches contain the novel mutations in Mendelian ratios, and are screened for recessive phenotypes of interest which appear in approximately 25% of embryos (Mullins et al., 1994). These embryos are referred to as “mutants” whereas those without phenotypes are “siblings”.

Mutant embryos are homozygous for a novel allele (the “causative mutation”) and due to genetic linkage, they are likely to be homozygous for alleles physically nearby on the chromosome. The location encompassing the causative mutation therefore lies in a region which is highly homozygous in mutants, yet heterozygous in siblings. This is referred to as linkage disequilibrium (LD). The region of high LD can be mapped using high-throughput sequencing and bioinformatics pipelines (Mackay & Schulte-Merker, 2014; Minevich et al., 2012; Obholzer et al., 2012) whereas prior efforts involved painstaking genotyping of simple sequence length polymorphisms and genome walks using bacterial or P1 artificial chromosome libraries or later microarrays (Stickney et al., 2002; Zhang et al., 1998).

All mapping processes rely on identification of polymorphic loci throughout the genome. Laboratory zebrafish strains are known to have a high degree of intra-strain polymorphism (Guryev et al., 2006), but mapping is aided by the introduction of alleles from other strains. Thus, mutagenised males are often paired with females from a different strain. As a result, alleles in the mutants and siblings are inherited from two different strains. This remains true throughout the multiple generations that a mutant line is maintained in a laboratory.

In this study, we report that the highly polymorphic nature of zebrafish strains can lead to gene expression differences between mutant and sibling embryos through allele-specific expression (ASE). The effect of ASE is well documented across many species, and can be tissue- and condition-specific (Ayroles et al., 2009; Doss et al., 2005; Fu et al., 2009; GTEx Consortium, 2017; Huang et al., 2015; Kim-Hellmuth et al., 2020; Storey et al., 2005). Here, this phenomenon manifests as a cluster of differentially expressed genes located near to the causative mutation site in many different unrelated mutant lines. Most often, the differential transcript levels of these local genes are likely due to expression differences between wild-type strains rather than altered transcription due to the mutation. We confirm the high prevalence of ASE in zebrafish in the SAT line which is derived from only two haplotypes. This observation has implications for researchers attempting to use differential expression to explain phenotypes of interest, not only in zebrafish, but also in other outbred model organisms, as these local genes may simply be a red herring.

## Results

### Differentially expressed genes are often enriched on the mutant chromosome

To map the causal mutations for a number of different mutants from forward genetics screens, we used RNA-seq and linkage disequilibrium (LD) mapping, based on Cloudmap (Minevich et al., 2012). A representative LD mapping plot (taken from the mutant line *u426*) is shown in Figure 1. We observed a high degree of LD on chromosome 7 at approximately 22 Mbp, suggesting the phenotype-causing mutation is near this position. DESeq2 reported 209 genes as differentially expressed (adjusted p-value<0.05) between mutants and siblings. Annotating the LD mapping plot with the position of these genes showed a cluster of differentially expressed (DE) genes near the LD mapping peak on chromosome 7. Indeed, we found 15 DE genes in an arbitrarily sized 20 Mbp window centred on the mapping peak at 22 Mbp, representing 7% of all DE genes. For comparison, a 20 Mbp window randomly sampled (1000 iterations) from the zebrafish genome contains approximately 1.4% of known genes.

**Figure 1:**
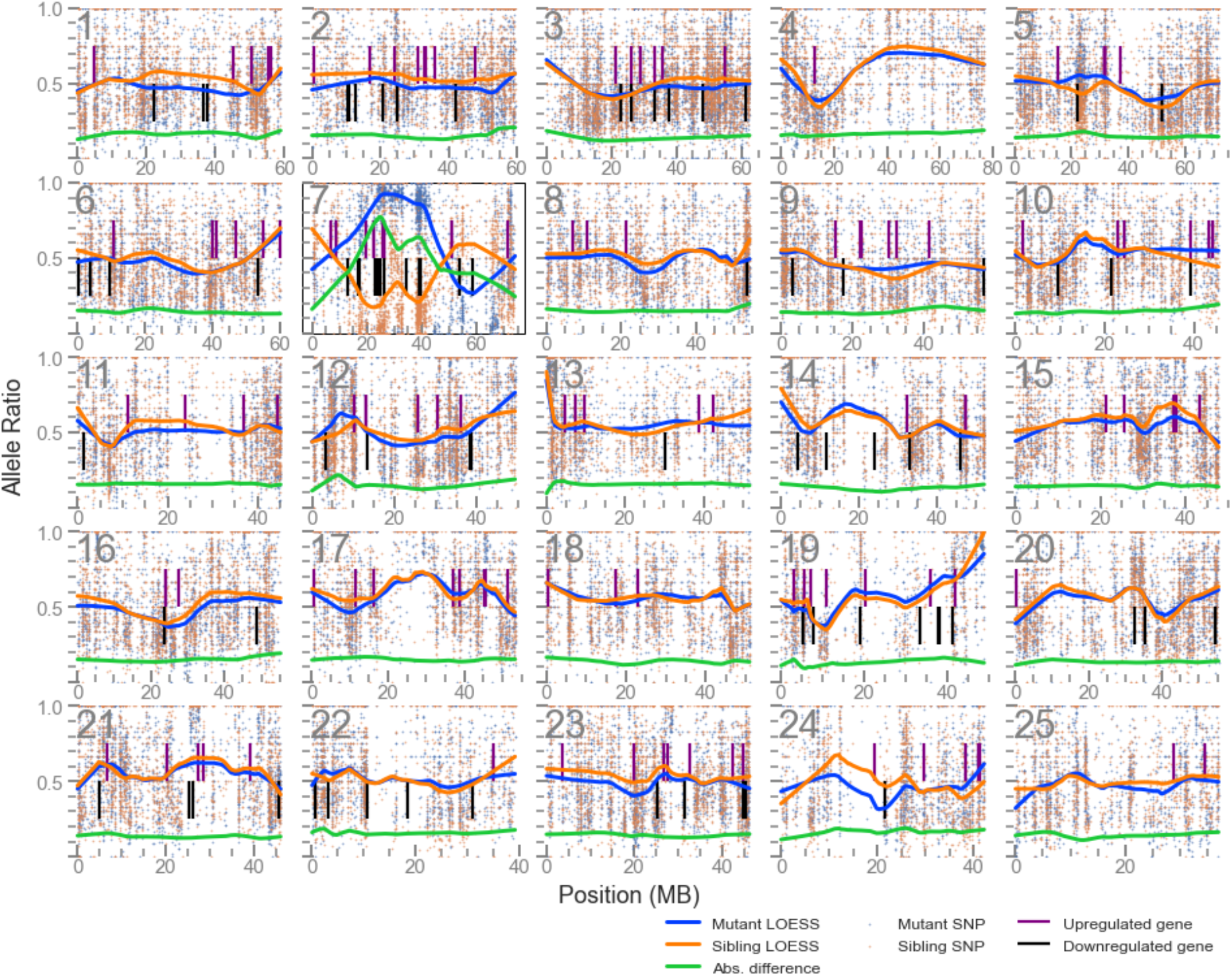
LD mapping plot of up and downregulated genes in *u426* mutants shows a cluster of such genes local to the mutation site on chromosome 7. The scatter plots for each of the 25 chromosomes shows the allele balance (proportion of reads containing the alternative allele) of each single nucleotide polymorphism (SNP) locus along with its physical position. Blue dots represent data from the mutant pool, and orange from the sibling pool. The blue and orange lines are LOESS-smoothed averages of these data and the green line is the absolute difference of the mutant and sibling samples, and is used to identify the region of highest LD. Vertical lines indicate the position of differentially expressed genes.

We then used a logistic regression model to examine the effect of LD on the probability of an individual gene being differentially expressed. A summary of each line and the regression results are presented in Table 1. Out of 9 mutant lines analysed, 7 samples showed a significant, positive effect of LD (Benjamini/Hochberg adjusted P value < 0.05). To help visualise the effect of LD on DE probability we calculated an odds ratio for each sample by comparing the DE probability at the site of maximum LD with the probability at a site of median LD. In the most extreme case (the sample *nl14*), the likelihood of finding a DE gene near to the mutation site is over 100-fold higher than the likelihood of finding one at a random other location in the genome.

**Table 1:** Summary of logistic regression results for RNA-seq analysed mutant lines. Causative mutation. shows the gene and location of the mutation site in lines where this has been confirmed empirically, otherwise the location is estimated from LD data. **Significance** column indicates adjusted p-value (***: <0.001, **: <0.01; *: <0.05). **Odds ratio** compares DE likelihood at maximum LD versus site of median LD. The **Nearby genes** column shows the number of DE genes lying within a 20Mbp window centred on the mutation site, and the percentage of these genes out of the total DE genes. In-table citations: ^1^(Barlow et al., 2020), ^2^(Miesfeld et al., 2015), ^3^(Armant et al., 2016). *nl14* line kindly provided by Alex Nechiporuk.

| Allele | Causative mutation | DE genes / total | Coefficient ± SEM | Sig. | Odds ratio | Nearby genes (%) |
| --- | --- | --- | --- | --- | --- | --- |
| *nl14* | *lama1* unpublished (chr24, 41.6Mbp) | 12 / 31664 | 9.09 ± 1.56 | *** | 118.5 | 3 (25%) |
| *la015577<sup>1</sup>* | *dmist* (chr5, 19.9Mbp) | 157 / 31199 | 6.84 ± 0.46 | *** | 55.8 | 23 (15%) |
| *u505<sup>1</sup>* | *dmist* (chr5, 19.9Mbp) | 71 / 31199 | 8.72 ± 0.72 | *** | 44.0 | 13 (18%) |
| *u757* | unpublished (chr23, 22Mbp) | 33 / 31199 | 6.31 ± 2.13 | ** | 7.8 | 1 (3%) |
| *u534* | not known (chr1, ~25Mbp) | 87 / 31664 | 4.83 ± 1.05 | *** | 5.4 | 4 (5%) |
| *u426* | not known (chr7, ~22Mbp) | 209 / 31664 | 2.67 ± 0.48 | *** | 5.3 | 15 (7%) |
| *nl13<sup>2</sup>* | *yap1* (chr18, 37.2Mbp) | 140 / 31199 | 2.58 ± 1.57 | - | 2.3 | 4 (3%) |
| *sb55<sup>3</sup>* | *ache* (chr 7, 26.0Mbp) | 348 / 24558 | 3.77 ± 1.67 | * | 2.0 | 14 (4%) |
| *u535* | not known (chr13, ~25Mbp) | 294 / 31663 | 0.35 ± 1.04 | - | 1.1 | 4 (1%) |

In parallel, we were analysing a separate catalogue of 3’ tag sequencing experiments of zebrafish mutant lines (115 experiments), most of which were generated and made available as part of the Zebrafish Mutation Project (Collins et al., 2015; Dooley et al., 2019; Kettleborough et al., 2013). These were analysed for differential expression producing a large collection of differentially expressed gene lists. We noticed that, often, the mutant chromosome had a large proportion of the total number of differentially expressed genes in the experiment. For example, comparing *mitfa*^*w2/w2*^ embryos to siblings produces 116 differentially expressed genes, 48 of which are present on chromosome 6, which is the chromosome where *mitfa* is located (Figure 2A).

**Figure 2.**
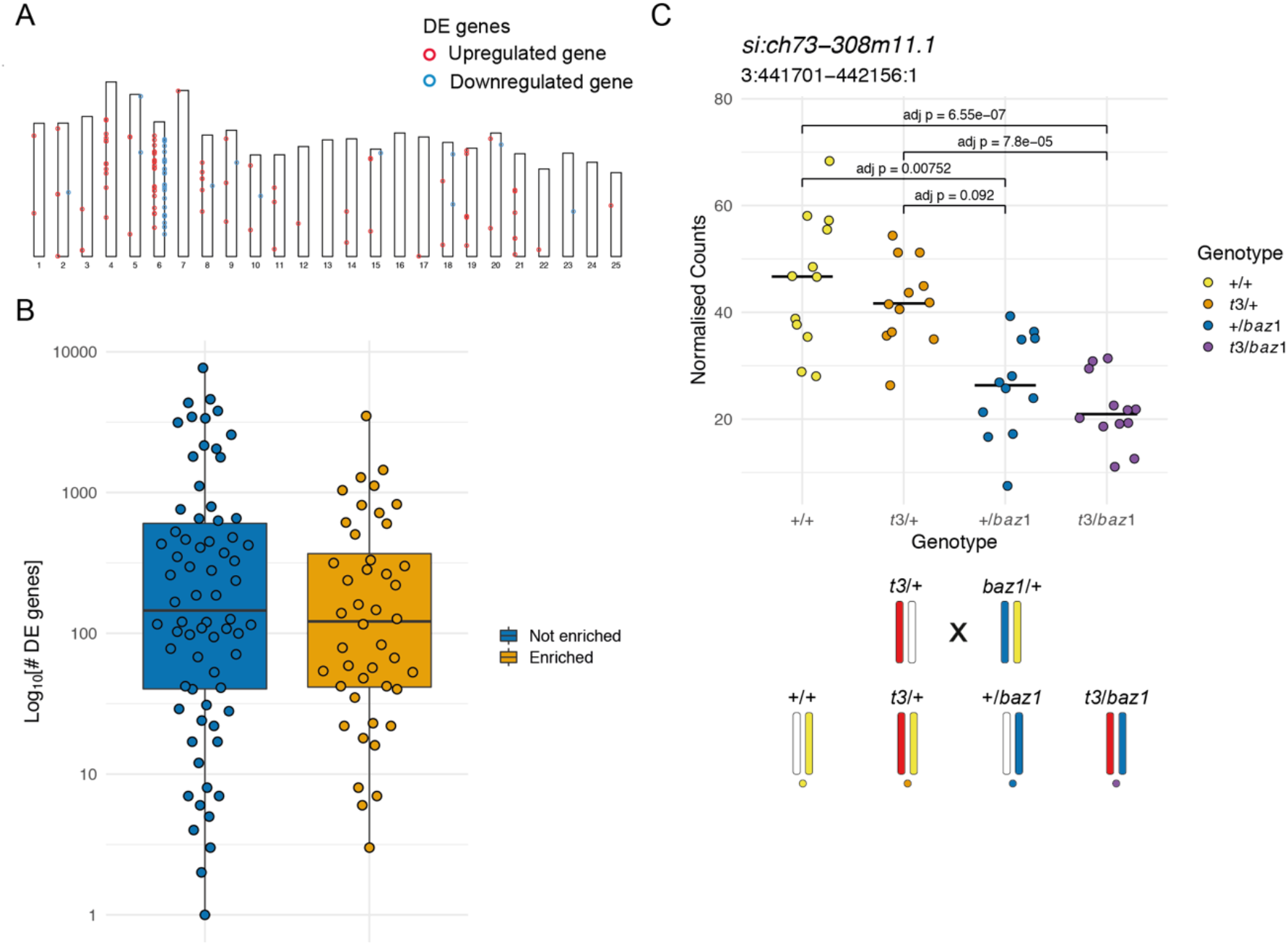
Enrichment of differentially expressed genes on the mutant chromosome. **A.** Ideogram showing the locations of the differentially expressed (DE) genes in a *mitfa*^*w2*^ incross. Circles represent differentially expressed genes and are coloured red if the gene is upregulated in the mutant embryos and blue if it is downregulated. **B**. Distribution of the total number of differentially expressed genes in experiments according to whether there is an enrichment on the mutant chromosome (orange) or not (blue), plotted on a log10 scale. **C**. Plot of normalised counts according to genotype in an intercross of two different *sox10* alleles. Yellow = wild type (+/+), orange = *sox10 t3* heterozygotes (*t3*/+), blue = *sox10 baz1* heterozygotes (+/*baz1*), purple = *sox10 t3, baz1* compound heterozygotes (*t3*/*baz1*). The schematic below the plot shows the chromosomes contributing to each genotype. Embryos that share the wild-type allele inherited from the *baz1*/+ parent (yellow chromosome) show higher expression levels.

To investigate this, we tested for chromosomes that had an enrichment of differentially expressed genes under the null hypothesis that they are randomly distributed across the genome. In all, 60 chromosomes from 37 lines had a statistically significant (binomial test, Bonferroni adjusted p < 0.05) enrichment of the DE genes. Of these, 44 were on the chromosome carrying the mutation being investigated in the experiment (Supplementary Table 1). Of the other 16, 7 had an enrichment on chromosome 9. This is driven by expression of γ-crystallin genes (Supplementary Table 2), which are expressed in the lens and present in a cluster on chromosome 9 (Greiling et al., 2009) that we have previously observed as being co-regulated (White et al., 2017). This suggests that the eyes are affected in some of the analysed mutants. Whether there is an enrichment of DE genes on the mutant chromosome does not depend on the total number of DE genes found in the experiment, although experiments with very high numbers of DE genes tend not to show an enrichment (Figure 2B).

In one experiment, we noticed that the differential expression of some genes was linked to one of the wild-type chromosomes in the experiment. This experiment was an intercross of two different *sox10* alleles, *t3* (Dutton et al., 2001) and *baz1* (Carney et al., 2006). Embryos were genotyped for both *sox10* alleles, which allowed us to also track the wild-type chromosomes in the cross. We noticed that two of the genotypes had expression levels for some genes on the same chromosome as *sox10* that were different from the other two genotypes (Figure 2C). The groups with higher expression share the wild-type chromosome from the *baz1*/+ parent (Figure 2C, yellow chromosome) whereas the others share the chromosome carrying the *baz1* allele (Figure 2C, blue chromosome). One explanation for this is that there is higher expression from the *si:ch73–308m11*.*1* allele on the wild-type chromosome (Figure 2C, yellow chromosome), which led us to hypothesise that the enrichment of differentially expressed genes on the mutant chromosome is not necessarily dependent on the mutant gene.

Our hypothesis is that allele-specific expression (ASE), i.e. polymorphism-driven variation in expression levels of genes, is common across the genome. This would manifest as differential expression when a genomic locus is driven to homozygosity in some individuals and the expression levels of genes in this locus are compared to those in individuals that are heterozygous, or homozygous for the other allele.

### Allele-specific expression is common in a wild-type cross

To test the hypothesis that the over-representation of differentially expressed genes on the mutant chromosome is driven by ASE and is independent of a mutated gene, we investigated gene expression in wild-type fish with defined haplotypes to enable easy identification of the different alleles in the cross. We used the SAT line, which was generated from an incross of one fully sequenced double haploid AB fish and one fully sequenced double haploid Tübingen fish (Howe et al., 2013).This means that for any position in the genome there are up to two possible alleles. The original haplotypes have recombined through the generations that the SAT line has been maintained by incrossing. We incrossed two SAT fish, fin-clipped them to isolate DNA for exome sequencing, collected 96 morphologically normal embryos at 5 days post fertilisation (dpf), extracted RNA from the individual embryos, and did RNA-seq on the 96 samples. We used the exome sequence of the SAT parent fish for this cross to call SNPs and identify regions that are either homozygous for the AB haplotype, homozygous for the Tübingen haplotype or heterozygous. Using the RNA-seq reads and SNPs identified in the parental exome data, we genotyped the embryos at locations that distinguish the AB and Tübingen haplotypes. Aggregating these data in 1 Mbp regions allowed us to determine the haplotypes of each individual embryo. We identified regions of the parental genomes where at least two genotypes, and thus potentially allele-specific expression, are possible in the offspring (informative regions) and where we had sufficient read depth to unambiguously identify the haplotypes in the offspring. We grouped the 96 RNA-seq samples according to their haplotype in that region (Figure 3A–B). Across the genome, this resulted in 82 different groupings of embryos according to local genotype. Embryos that had evidence of a recombination event within the informative region were assigned to a genotype group according to the largest contiguous section of the region.

**Figure 3.**
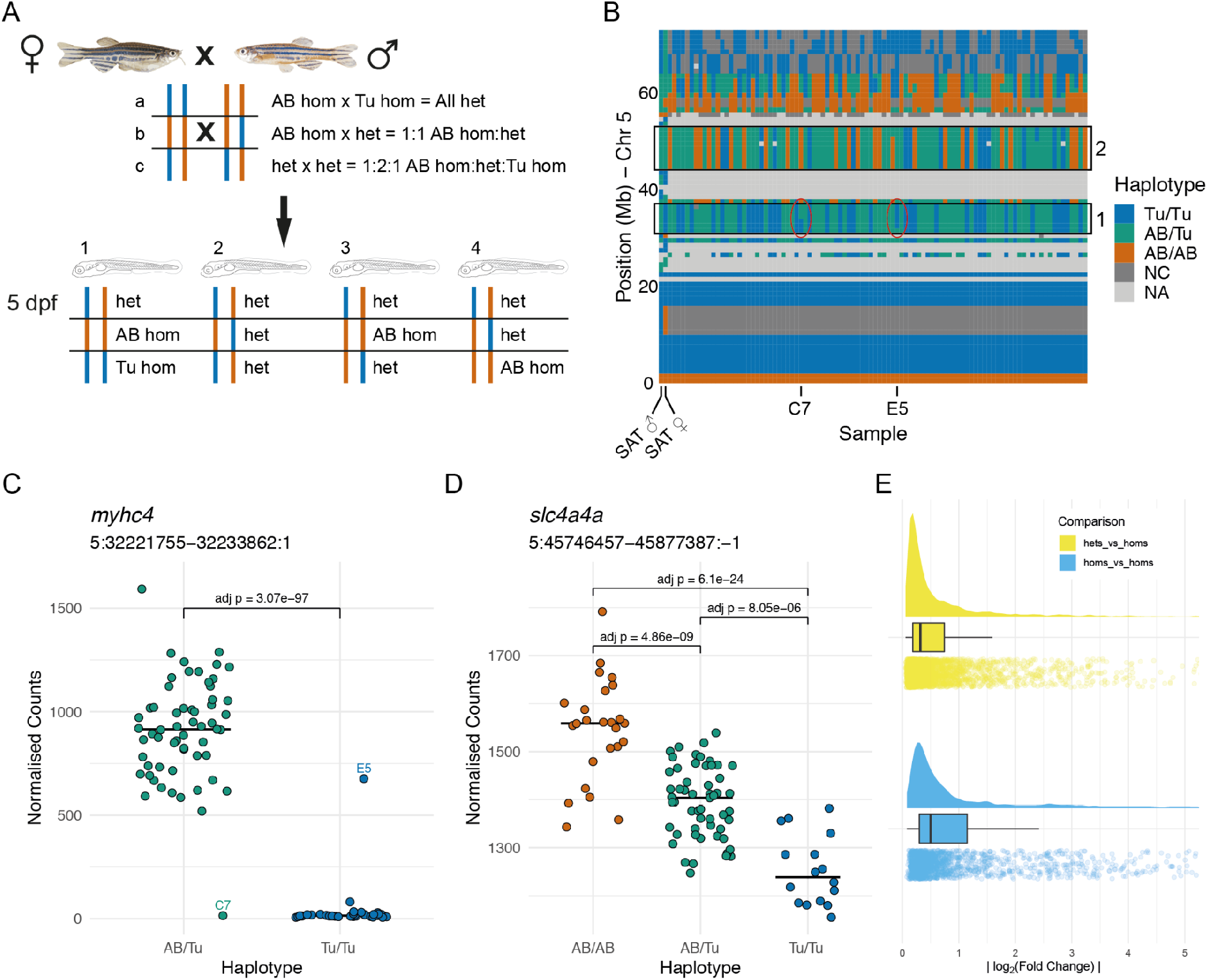
Allele-specific expression is common in wild-type embryos. **A**. Experimental design. Two wild-type SAT fish were incrossed and 96 embryos were collected for RNA-seq at 5 dpf. Depending on the haplotypes of the parents, different combinations of genotype are possible in specific regions in the offspring. **B**. The haplotypes of the collected embryos were determined in 1 Mbp bins using the RNA-seq reads and the embryos were grouped according to the haplotypes in specific regions. Chromosome 5 is shown with samples along the x-axis and chromosomal position on the y-axis. 1 Mbp bins are coloured according to the haplotype in that region. Blue = homozygous Tübingen (Tu/Tu), green = heterozygous AB/Tübingen (AB/Tu), orange = homozygous AB (AB/AB), dark grey = Not consistent with parental haplotypes (NC), light grey = No haplotype call (NA), due to, for example, low coverage. Examples of regions used to group the embryos are boxed. **C–D**. Examples of differentially expressed genes from two different groupings. **C**. Counts for the *myhc4* gene, grouped according to the haplotypes in the region 5:31–37 Mbp (region 1 in B). The Tübingen allele is expressed at very low levels, with much higher expression in the heterozygotes. There are two examples of embryos with recombinations within the region. Compare to circled areas in the haplotype plot in B. **D**. Example of a differentially expressed gene (*slc4a4a*) in a region where all three genotypes are present (5:44–53 Mbp, region 2 in B). As in C, the Tübingen allele has lower expression, with the heterozygotes showing intermediate levels. **E**. Distribution of absolute log_2_(fold change) values found between wild-type alleles. Differences when comparing homozygous embryos (blue) are generally larger than when comparing heterozygotes to homozygotes (yellow).

We then ran differential gene expression analysis on each different embryo grouping, which showed that differentially expressed genes were located in or close to the region of the genome that was used to define the embryo groups (Figure 3 and Supplementary Figure 1, Supplementary Table 3). The log_2_(fold changes) of affected genes varied widely, but had an absolute mean of 0.5 for the homozygous versus homozygous comparison (Figure 3E). This demonstrates that genes show allele-specific expression in a wild-type context (Figure 3C– E).

Interestingly, it is possible to see the consequences of meiotic recombination in individual embryos (Fig 3B–C). For example, two samples (C7 and E5) show recombination in the 31– 37 Mbp region of chromosome 5 (red ovals in Figure 3B). The genotype near the *myhc4* gene is actually the opposite of that called for the whole region and this is evident in the count plot – C7 has expression comparable with the Tu/Tu haplotype, despite being assigned AB/Tu and E5 has expression similar to the AB/Tu sample despite being assigned Tu/Tu based on the entire 31–37 Mbp region (Figure 3C).

**Supplementary Figure 1.**
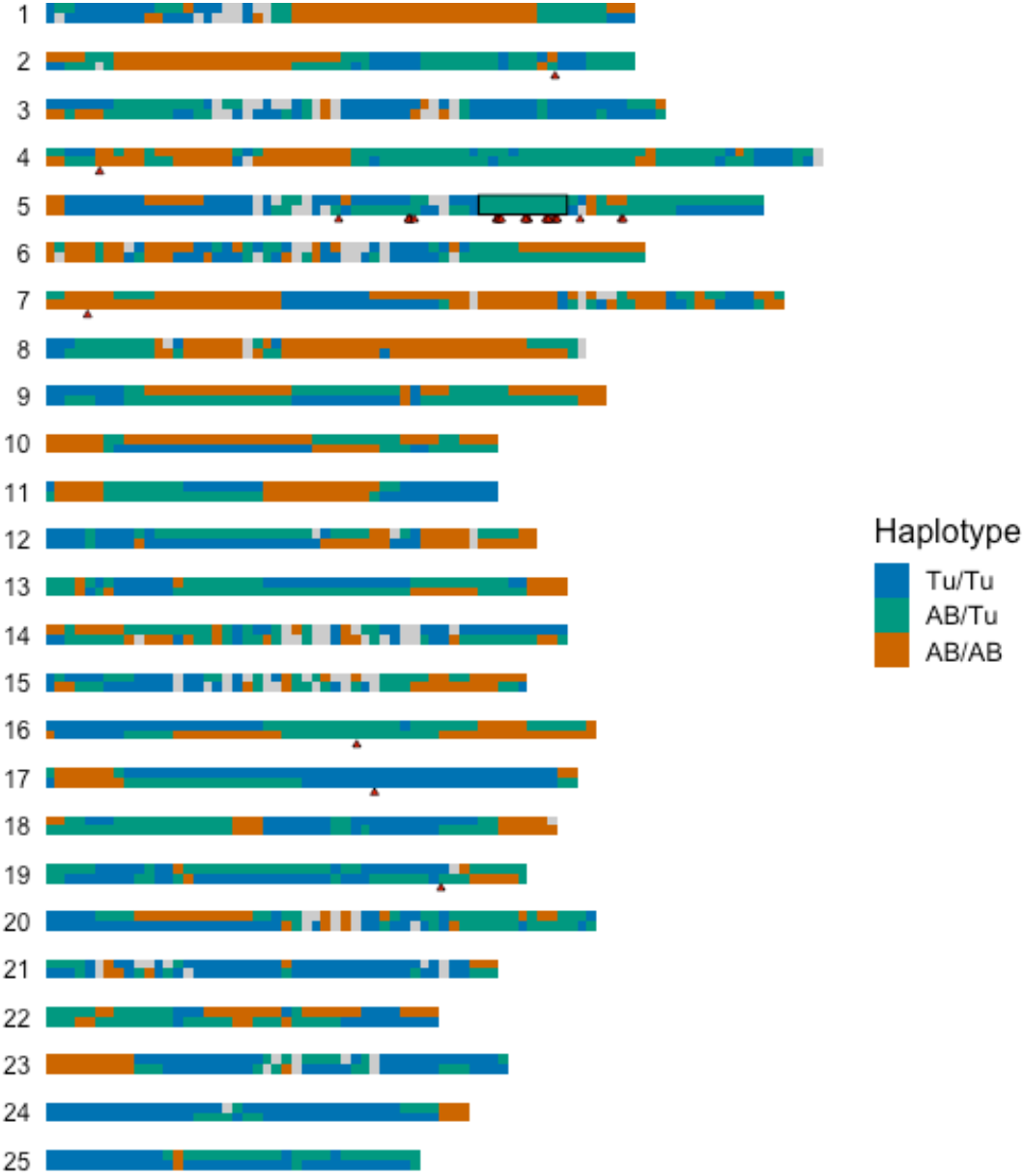
Allele-specific expression is linked to the region used to define the sample groupings. Representation of the parental haplotypes of the SAT cross across all 25 chromosomes (blue = Tu/Tu, green = AB/Tu, orange = AB/AB). The box shows the region (Chr5:44-53 Mb) that was used to define the groups of embryos compared using DESeq2. The red triangles show the positions of the differentially expressed genes, most of which are in or close to the region.

### Distinguishing response genes from allele-specific expression

Having established that ASE is widespread and can significantly alter the transcriptional profiles of mutant zebrafish, we wondered whether there is a way to distinguish potential “true” response genes located on the same chromosome as the mutation, i.e. those that change expression due to the altered function of the mutated gene, from those differentially expressed genes that arise through ASE. We went back to the expression data from the compound heterozygous *sox10*^*t3*^;*sox10*^*baz1*^ cross and found that the genes that were differentially expressed between *sox10*^*t3/baz1*^ individuals and their siblings and located on chr3 fall into different groups with respect to their expression levels across the four different genotypes (Figure 4). Ten genes showed expression patterns as shown in Figure 2C, where increased expression is linked to the presence of a specific allele (Figure 4A). Only one gene (ENSDARG00000110416) located on another chromosome, encoding a miRNA, showed a similar pattern (Supplementary Figure 2). By contrast, the other 15 differentially expressed genes (excluding *sox10* itself) on chr3 showed genotype-dependent transcript levels that were consistent with (though do not prove) a response to loss of *sox10* function, i.e. the wild types and the compound heterozygous individuals had opposing expression levels whereas both heterozygous genotypes had intermediate levels or the same as wild types (Figure 4B). However, the genes showing single allele-linked expression patterns suggest that ASE is the primary driver of their differential expression and that they are probably red herrings.

**Figure 4.**
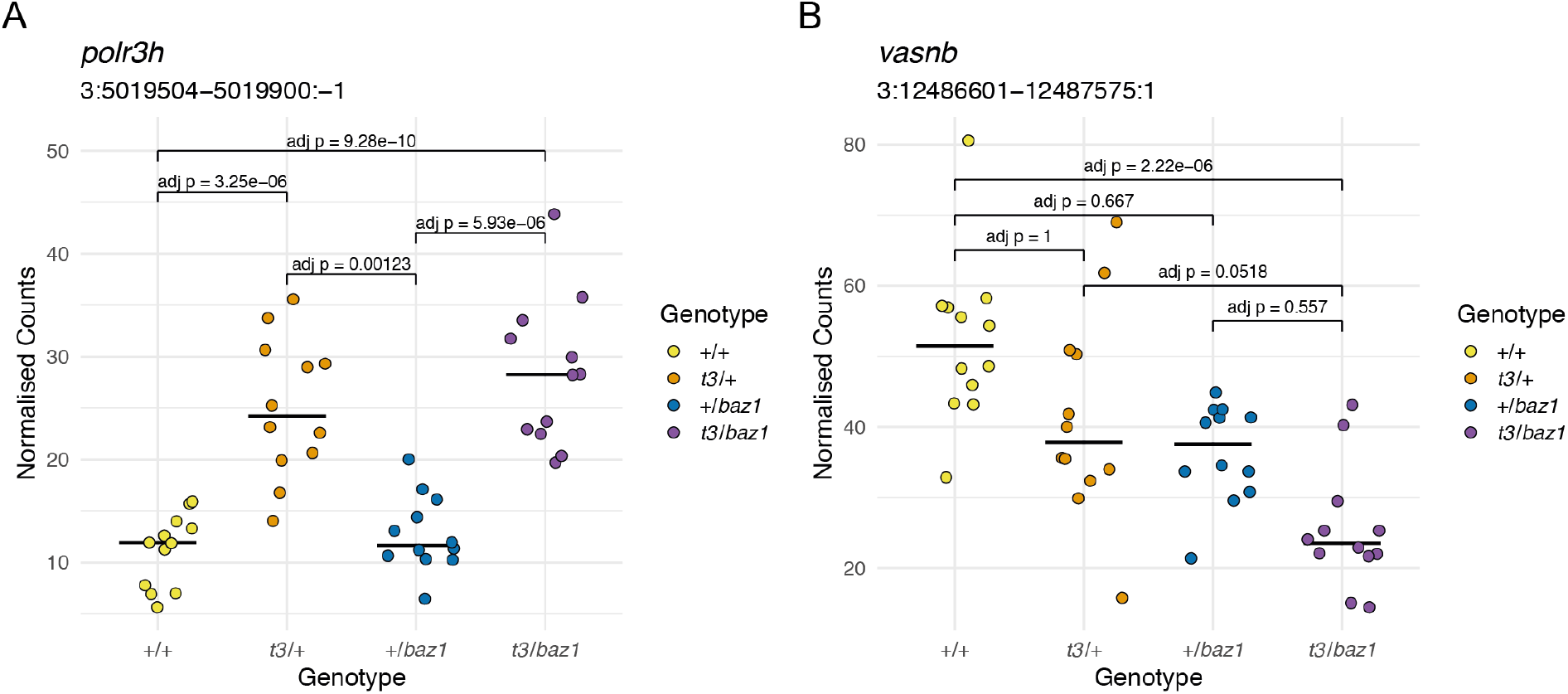
Distinguishing mutation-dependent gene expression changes from allele-specific expression. **A**. Plot of normalised counts consistent with ASE. This shows either reduced expression from the allele on one of the wild-type chromosomes (white chromosome in diagram in Figure 2C) or increased expression from the allele on the *t3* chromosome. **B**. Normalised counts consistent with a response to the *sox10* mutations. The compound heterozygotes have reduced expression and the other two groups of heterozygotes are intermediate between the compound heterozygotes and the wild-types. Yellow = wild-types (+/+), orange = *t3* heterozygotes (*t3*/+), blue = *baz1* heterozygotes (+/*baz1*), purple = compound heterozygotes (*t3*/*baz1*).

**Supplementary Figure 2.**
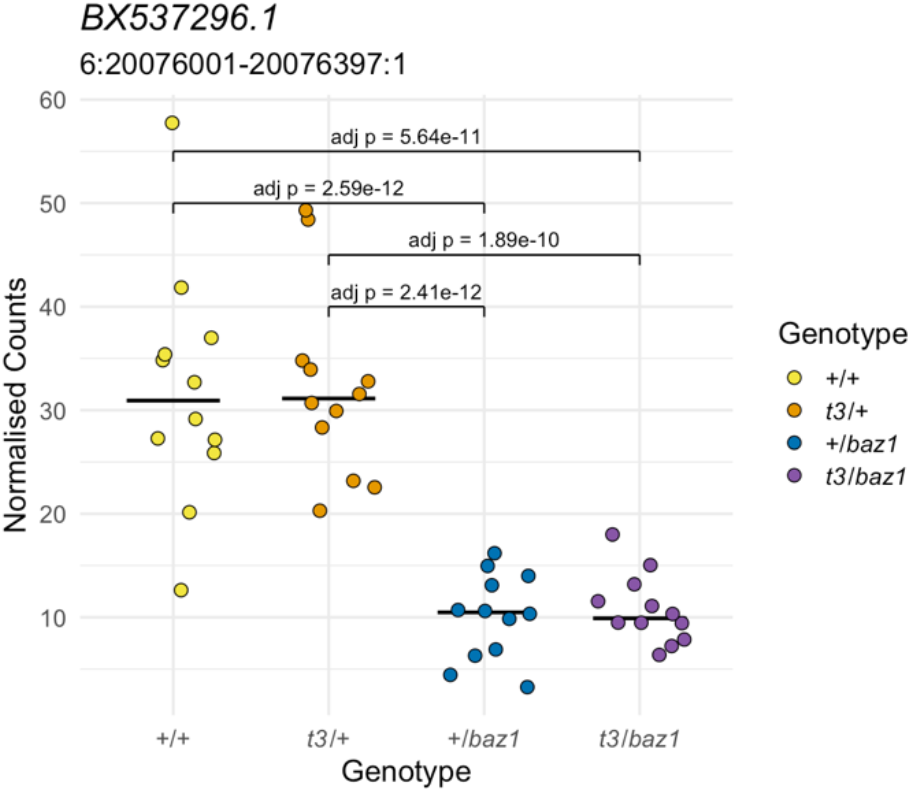
Allele-specific expression not linked to the homozygous region. The expression of this gene according to genotype is not consistent with a response to a recessive mutation. Yellow = wild-types (+/+), orange = *t3* heterozygotes (*t3*/+), blue = *baz1* heterozygotes (+/*baz1*), purple = compound heterozygotes (*t3*/*baz1*).

## Discussion

Transcriptional profiling is a powerful and popular technique to investigate the gene expression changes resulting from organismal insults such as drug treatments, infections or altered gene function. To gain mechanistic insight into gene regulatory events affected by a particular mutation, it is paramount to distinguish specific responses due to altered function of the mutated genes from other causes that change transcript abundance, such as developmental delay or technical artefacts such as batch effects. In this work, we describe a previously under-appreciated effect of allele-specific expression on the transcriptomes of zebrafish mutants. In 51 out of 124 transcriptional profiling experiments comparing zebrafish mutants and siblings at different stages of development, we found a statistically significant enrichment of differentially expressed genes on the same chromosome as the mutated gene.

The physical arrangement of genes in an organism’s genome is not random. Co-expression of functionally related genes using shared regulatory elements and/or transcription factors provides an evolutionary pressure to keep these genes clustered in physical proximity within a chromosome (Thévenin et al., 2014). Consequently, it is possible that a mutation affecting one gene could alter expression levels of nearby genes if they form a functionally related cluster. However, this was not the case for the neighbouring differentially expressed genes in the tested mutant lines. (Of note, 7/16 chromosomal enrichments that were not linked to the mutated genes affected a chromosome 9 cluster of crystallin genes that are expressed in the eye.) Alternatively, differential transcript abundance between homozygous mutants and siblings could arise because different transcript levels are produced from the stretch of homozygous genome surrounding the mutation than from the corresponding genomic region in the siblings, which are heterozygous or homozygous for the other allele. This allele-specific expression has been widely demonstrated across different tissues and organisms (GTEx Consortium, 2017; Huang et al., 2015; Kim-Hellmuth et al., 2020) and can play a role in developmental and disease processes (Libioulle et al., 2007; Moffatt et al., 2007; Nicolae et al., 2010).

We tested the contribution of allele-specific expression using RNA-seq in the zebrafish SAT wild-type strain that was generated from two fish double haploid for either the Tübingen background or the AB background. We found that we could elicit the same effect of differential expression clustering when we compared individuals homozygous for one haplotype with their siblings that were heterozygous or homozygous for the other haplotype. In both cases, due to recombination in the parents of a cross, in a comparison of homozygous mutants and their siblings, this effect is greatest closest to the mutation. Indeed, we can relate recombinations observed in the haplotype calls to expression levels in individual embryos. ASE is often tissue-dependent and the average log2(fold change) between alleles in human ASE is about 0.6 as measured in different tissues (GTEx Consortium, 2017). Here we can observe ASE at similar magnitudes even when averaged across all tissues through whole embryo RNA-seq. This suggests that the expression differences between alleles would be even larger when looking at individual tissues.

Zebrafish wild-type “strains” are not strains in the same sense as the well-characterised inbred lines in mouse or medaka, for example. Zebrafish are highly polymorphic, such that allele-specific expression is evident even in lines that were not outcrossed to another genetic background before the experiment. This effect would be expected to be much less pronounced in RNA-seq data from inbred mouse strains. Indeed, in our work on RNA-seq data from mouse knockouts (Collins et al., 2019), we did not observe enrichment of differentially expressed genes on the mutant chromosome. However, allele-specific expression needs to be considered when working either in wild mouse strains, crosses between different genetic backgrounds, or indeed any organism that isn’t fully inbred.

Given that ASE can lead to differential expression between mutants and siblings, can we correct for it in transcript profiling experiments? The solution is not as simple as removing any differentially expressed genes in the same region of the chromosome as the mutation being studied. This is because the differentially expressed genes on the same chromosome as the mutation are likely to be a mix of genuine responses to the mutation and linkage of ASE unrelated to the mutation. One way to resolve this would be to use two different mutant alleles of the same gene to generate compound heterozygotes and enable tracking of parental alleles. This would allow genotyping for both alleles and the ability therefore to also identify the different wild-type chromosomes in the cross. As shown in Figure 4, this makes it possible to distinguish between potential genuine responses to the mutation and spurious ones. Another possibility would be to identify an informative SNP in the wild-type alleles of the mutant gene being studied to allow genotyping of both the mutation and the wild-type alleles. Both of these solutions would involve considerable effort and expense, and so would need careful consideration with respect to the need to validate specific gene expression changes for the conclusions of the study. As a first step, we recommend that, whatever analysis pipeline is used, the output of differentially expressed genes contains the locations of the genes, making it possible to easily see which genes are on the same chromosome as the mutation.

## Materials and Methods

### RNA-seq and LD mapping

Eight independent mutant fish lines under study by groups at UCL (zebrafishucl.org) were analysed by RNA-seq in order to simultaneously gain gene expression data and to measure alleles across the genome in order to help map the causative mutation. Seven of these lines were the product of ENU random mutagenesis, one was created by a random viral insertion, and one by a targeted CRISPR insertion. An additional sample was taken from the literature (Armant et al., 2016) at random by searching Pubmed for papers where RNA-seq data had been uploaded to the European Nucleotide Archive.

In preparation for RNA-seq, embryos or larvae were sorted into two groups based on their phenotypes (mutant and sibling), each comprising three pools of at least 15 individuals. RNA was extracted from these six samples and sequenced using the IIlumina NextSeq platform (2×75bp reads, approximately 75 million reads per sample). Reads were aligned to the GRCz10 genome using HISAT2 (Kim et al., 2019). To measure differential expression, transcripts were counted from the aligned RNA-seq reads using featureCounts (Liao et al., 2014) and compared using DESeq2 (Love et al., 2014). A gene was considered differentially expressed if the adjusted p-value from DESeq2 was below 0.05.

To perform LD mapping, the three samples in each group were analysed as a single pooled sample for single nucleotide polymorphisms (SNPs) by BCFtools (Li, 2011), calculating the allele ratio at each SNP location. SNPs which appeared in only one of the two genotype pools were filtered out, as were those with a quality score below 100. The absolute difference between a given SNP’s mutant and sibling allele ratio indicates the degree of segregation of that allele (Mackay & Schulte-Merker, 2014). These values can be smoothed using LOESS, producing maps of the genome showing regions of high LD (Minevich et al., 2012). The physical location of each gene’s start codon in the GRCz10 genome assembly was downloaded from Ensembl BioMart and appended to the DESeq2 table. The LD value was estimated at each gene’s position based on interpolation of the LOESS-smoothed SNP data. Finally, a logistic regression model was used to test the effect of LD on a gene’s probability of being differentially expressed. This was performed using the Logit function of the Python module statsmodels.

### DeTCT sequencing

DeTCT libraries were generated, sequenced and analysed as described previously (Collins et al., 2015). The resulting genomic regions and putative 3′ ends were filtered using DeTCT’s filter_output script (https://github.com/iansealy/DETCT/blob/master/script/filter_output.pl) in its --strict mode. -- strict mode removes 3’ ends in coding sequence, transposons, if nearby sequence is enriched for As or if not near a primary hexamer. Regions not associated with 3′ ends are also removed. Differential expression analysis was done using DeTCT’s run_pipeline script (https://github.com/iansealy/DETCT/blob/master/script/run_pipeline.pl) using DESeq2 (Love et al., 2014) with an adjusted p value cut-off of 0.05. Sequence data were deposited in the European Nucleotide Archive (ENA) under accessions ERP001656, ERP004581, ERP006132, ERP003802, ERP004579, ERP005517, ERP008771, ERP005564, ERP009868, ERP006133, ERP009078 and ERP013835.

### DNA sequencing

Double haploid AB and Tübingen fish were produced and sequenced as described in (Howe et al., 2013). Whole genome sequencing data (SRA Study: ERP000232) was downloaded from the European Nucleotide Archive. Exome sequencing on parents for the wild-type SAT cross was done as described (Kettleborough et al., 2013). Reads were mapped to the GRCz11 genome assembly using BWA (Li & Durbin, 2010, v0.5.10) and duplicates were marked with biobambam (Tischler & Leonard, 2014). SNPs were called using a modified version of the 1000 Genomes Project variant calling pipeline (The 1000 Genomes Project Consortium, 2010). Initial calls were done by SAMtools mpileup (Li, 2011), QCALL (Le & Durbin, 2011) and the GATK Unified Genotyper (DePristo et al., 2011). SNPs not called by all three callers were removed from the analysis, along with any SNP that did not pass a caller’s standard filters. Additionally, SNPs were removed where the genotype quality was lower than 100 for GATK and lower than 50 for QCALL and SAMtools mpileup and where the mean read depth per sample was less than 10. These SNP calls were then filtered for positions that are informative of the parental background in the SAT cross, i.e. ones that are homozygous reference in one double haploid fish and homozygous alternate in the other.

### RNA-seq of wild-type SAT embryos

RNA was extracted from 5 days post fertilisation (dpf) larvae as described previously (Wali et al., 2021). Briefly, RNA was extracted from individual embryos by mechanical lysis in RLT buffer (Qiagen) containing 1 μl of 14.3M β-mercaptoethanol (Sigma). The lysate was combined with 1.8 volumes of Agencourt RNAClean XP (Beckman Coulter) beads and allowed to bind for 10 minutes. The plate was applied to a plate magnet (Invitrogen) until the solution cleared and the supernatant was removed without disturbing the beads. This was followed by washing the beads three times with 70% ethanol. After the last wash, the pellet was allowed to air dry for 10 minutes and then resuspended in 50 μl of RNAse-free water. RNA was eluted from the beads by applying the plate to the magnetic rack. Samples were DNase-I treated to remove genomic DNA. RNA was quantified using Quant-IT RNA assay (Invitrogen). Stranded RNA-seq libraries were constructed using the Illumina TruSeq Stranded RNA protocol after treatment with Ribozero. Libraries were pooled and sequenced on 6 Illumina HiSeq 2500 lanes in 75 bp paired-end mode. Sequence data were deposited in ENA under accession ERP011556. Reads for each sample were aggregated across lanes (median reads per embryo = 18.1 M) and mapped to the GRCz11 zebrafish genome assembly using TopHat (Kim et al., 2013, v2.0.13, options: --library-type fr-firststrand). The data were assessed for technical quality (GC-content, insert size, proper pairs, etc…) using QoRTs (Hartley & Mullikin, 2015). Counts for genes were produced using htseq-count (Anders et al., 2015, v0.6.0 options: --stranded=reverse) with the Ensembl v97 annotation as a reference. Differential expression analysis was done in R (R Core Team, 2019) with DESeq2 (Love et al., 2014) using a cut-off for adjusted p-values of 0.05. The samples were genotyped at the positions that were determined to be informative using the double haploid sequence using GATK’s SplitNCigarReads tool followed by the HaplotypeCaller (Poplin et al., 2017) on the RNA-seq data. The genotype calls were converted to their strain of origin (either DHAB or DHTu) and haplotypes were called by taking the most frequent genotype call in 1 Mbp windows. Any haplotypes that were not consistent with the parental haplotypes were removed.

## Data availability

Sequencing data have been deposited in ENA under the accessions shown in the Materials and Methods. Differentially expressed gene lists for all the experiments are available at doi.org/10.6084/m9.figshare.15082239.

## Acknowledgements

We thank G. Gestri, J. Rihel and T. Hawkins for providing RNA-seq datasets from mutant lines they are working on, Alex Nechiporuk for providing mutant lines, technical staff in our fish facilities for animal care and Ian Sealy and Munise Merteroglu for critical reading of the manuscript. This study was supported by MRC Programme Grant support to Gaia Gestri and SW (MR/L003775/1 and MR/T020164/1), and a Wellcome Trust Investigator Award to SW (095722/Z/11/Z).

## References

Anders, S., Pyl, P. T., & Huber, W. (2015). HTSeq—A Python framework to work with high-throughput sequencing data. Bioinformatics, 31(2), 166–169. https://doi.org/10.1093/bioinformatics/btu638

Armant, O., Gourain, V., Etard, C., & Strähle, U. (2016). Whole transcriptome data analysis of zebrafish mutants affecting muscle development. Data in Brief, 8, 61–68. https://doi.org/10.1016/j.dib.2016.05.007

Ayroles, J. F., Carbone, M. A., Stone, E. A., Jordan, K. W., Lyman, R. F., Magwire, M. M., Rollmann, S. M., Duncan, L. H., Lawrence, F., Anholt, R. R. H., & Mackay, T. F. C. (2009). Systems genetics of complex traits in Drosophila melanogaster. Nature Genetics, 41(3), 299–307. https://doi.org/10.1038/ng.332

Barlow, I. L., Mackay, E., Wheater, E., Goel, A., Lim, S., Zimmerman, S., Woods, I., Prober, D. A., & Rihel, J. (2020). A genetic screen identifies dreammist as a regulator of sleep. BioRxiv, 2020.11.18.388736. https://doi.org/10.1101/2020.11.18.388736

Carney, T. J., Dutton, K. A., Greenhill, E., Delfino-Machín, M., Dufourcq, P., Blader, P., & Kelsh, R. N. (2006). A direct role for Sox10 in specification of neural crest-derived sensory neurons. Development, 133(23), 4619–4630. https://doi.org/10.1242/dev.02668

Collins, J. E., Wali, N., Sealy, I. M., Morris, J. A., White, R. J., Leonard, S. R., Jackson, D. K., Jones, M. C., Smerdon, N. C., Zamora, J., Dooley, C. M., Carruthers, S. N., Barrett, J. C., Stemple, D. L., & Busch-Nentwich, E. M. (2015). High-throughput and quantitative genome-wide messenger RNA sequencing for molecular phenotyping. BMC Genomics, 16(1), 578. https://doi.org/10.1186/s12864-015-1788-6

Collins, J. E., White, R. J., Staudt, N., Sealy, I. M., Packham, I., Wali, N., Tudor, C., Mazzeo, C., Green, A., Siragher, E., Ryder, E., White, J. K., Papatheodoru, I., Tang, A., Füllgrabe, A., Billis, K., Geyer, S. H., Weninger, W. J., Galli, A.,… Busch-Nentwich, E. M. (2019). Common and distinct transcriptional signatures of mammalian embryonic lethality. Nature Communications, 10(1), 2792. https://doi.org/10.1038/s41467-019-10642-x

DePristo, M. A., Banks, E., Poplin, R., Garimella, K. V., Maguire, J. R., Hartl, C., Philippakis, A. A., del Angel, G., Rivas, M. A., Hanna, M., McKenna, A., Fennell, T. J., Kernytsky, A. M., Sivachenko, A. Y., Cibulskis, K., Gabriel, S. B., Altshuler, D., & Daly, M. J. (2011). A framework for variation discovery and genotyping using next-generation DNA sequencing data. Nature Genetics, 43(5), 491–498. https://doi.org/10.1038/ng.806

Dooley, C. M., Wali, N., Sealy, I. M., White, R. J., Stemple, D. L., Collins, J. E., & Busch-Nentwich, E. M. (2019). The gene regulatory basis of genetic compensation during neural crest induction. PLOS Genetics, 15(6), e1008213. https://doi.org/10.1371/journal.pgen.1008213

Doss, S., Schadt, E. E., Drake, T. A., & Lusis, A. J. (2005). Cis-acting expression quantitative trait loci in mice. Genome Research, 15(5), 681–691. https://doi.org/10.1101/gr.3216905

Driever, W., Solnica-Krezel, L., Schier, A. F., Neuhauss, S. C., Malicki, J., Stemple, D. L., Stainier, D. Y., Zwartkruis, F., Abdelilah, S., Rangini, Z., Belak, J., & Boggs, C. (1996). A genetic screen for mutations affecting embryogenesis in zebrafish. Development (Cambridge, England), 123, 37–46.

Dutton, K. A., Pauliny, A., Lopes, S. S., Elworthy, S., Carney, T. J., Rauch, J., Geisler, R., Haffter, P., & Kelsh, R. N. (2001). Zebrafish colourless encodes sox10 and specifies non-ectomesenchymal neural crest fates. Development (Cambridge, England), 128(21), 4113–4125. https://doi.org/10.1242/dev.128.21.4113

Fu, J., Keurentjes, J. J. B., Bouwmeester, H., America, T., Verstappen, F. W. A., Ward, J. L., Beale, M. H., de Vos, R. C. H., Dijkstra, M., Scheltema, R. A., Johannes, F., Koornneef, M., Vreugdenhil, D., Breitling, R., & Jansen, R. C. (2009). System-wide molecular evidence for phenotypic buffering in Arabidopsis. Nature Genetics, 41(2), 166–167. https://doi.org/10.1038/ng.308

Greiling, T. M. S., Houck, S. A., & Clark, J. I. (2009). The zebrafish lens proteome during development and aging. Molecular Vision, 15, 2313–2325.

GTEx Consortium. (2017). Genetic effects on gene expression across human tissues. Nature, 550(7675), 204–213. https://doi.org/10.1038/nature24277

Guryev, V., Koudijs, M. J., Berezikov, E., Johnson, S. L., Plasterk, R. H. A.Eeden, F. J. M. van, & Cuppen, E. (2006). Genetic variation in the zebrafish. Genome Research, 16(4), 491–497. https://doi.org/10.1101/gr.4791006

Haffter, P., Granato, M., Brand, M., Mullins, M. C., Hammerschmidt, M., Kane, D. A., Odenthal, J., van Eeden, F. J., Jiang, Y. J., Heisenberg, C. P., Kelsh, R. N., Furutani-Seiki, M., Vogelsang, E., Beuchle, D., Schach, U., Fabian, C., & Nüsslein-Volhard, C. (1996). The identification of genes with unique and essential functions in the development of the zebrafish, Danio rerio. Development (Cambridge, England), 123, 1–36.

Hartley, S. W., & Mullikin, J. C. (2015). QoRTs: A comprehensive toolset for quality control and data processing of RNA-Seq experiments. BMC Bioinformatics, 16(1), 224. https://doi.org/10.1186/s12859-015-0670-5

Howe, K., Clark, M. D., Torroja, C. F., Torrance, J., Berthelot, C., Muffato, M., Collins, J. E., Humphray, S., McLaren, K., Matthews, L., McLaren, S., Sealy, I., Caccamo, M., Churcher, C., Scott, C., Barrett, J. C., Koch, R., Rauch, G.-J., White, S.,… Stemple, D. L. (2013). The zebrafish reference genome sequence and its relationship to the human genome. Nature, 496(7446), 498–503. https://doi.org/10.1038/nature12111

Huang, W., Carbone, M. A., Magwire, M. M., Peiffer, J. A., Lyman, R. F., Stone, E. A., Anholt, R. R. H., & Mackay, T. F. C. (2015). Genetic basis of transcriptome diversity in *Drosophila melanogaster*. Proceedings of the National Academy of Sciences, 112(44), E6010. https://doi.org/10.1073/pnas.1519159112

Kettleborough, R. N. W., Bruijn E. de, Eeden, F. van, Cuppen, E., & Stemple, D. L. (2011). Chapter 6—High-Throughput Target-Selected Gene Inactivation in Zebrafish. In H. W. Detrich, M. Westerfield, & L. I. Zon (Eds.), Methods in Cell Biology (Vol. 104, pp. 121–127). Academic Press. https://doi.org/10.1016/B978-0-12-374814-0.00006-9

Kettleborough, R. N. W., Busch-Nentwich, E. M., Harvey, S. A., Dooley, C. M., de Bruijn, E., van Eeden, F., Sealy, I., White, R. J., Herd, C., Nijman, I. J., Fényes, F., Mehroke, S., Scahill, C., Gibbons, R., Wali, N., Carruthers, S., Hall, A., Yen, J., Cuppen, E., & Stemple, D. L. (2013). A systematic genome-wide analysis of zebrafish protein-coding gene function. Nature, 496(7446), 494–497. https://doi.org/10.1038/nature11992

Kim, D., Paggi, J. M., Park, C., Bennett, C., & Salzberg, S. L. (2019). Graph-based genome alignment and genotyping with HISAT2 and HISAT-genotype. Nature Biotechnology, 37(8), 907–915. https://doi.org/10.1038/s41587-019-0201-4

Kim, D., Pertea, G., Trapnell, C., Pimentel, H., Kelley, R., & Salzberg, S. L. (2013). TopHat2: Accurate alignment of transcriptomes in the presence of insertions, deletions and gene fusions. Genome Biology, 14(4), R36. https://doi.org/10.1186/gb-2013-14-4-r36

Kim-Hellmuth, S., Aguet, F., Oliva, M., Muñoz-Aguirre, M., Kasela, S., Wucher, V., Castel, S. E., Hamel, A. R., Viñuela, A., Roberts, A. L., Mangul, S., Wen, X., Wang, G., Barbeira, A. N., Garrido-Martín, D., Nadel, B. B., Zou, Y., Bonazzola, R., Quan, J.,… Lappalainen, T. (2020). Cell type–specific genetic regulation of gene expression across human tissues. Science, 369(6509), eaaz8528. https://doi.org/10.1126/science.aaz8528

Le, S. Q., & Durbin, R. (2011). SNP detection and genotyping from low-coverage sequencing data on multiple diploid samples. Genome Research, 21(6), 952–960. https://doi.org/10.1101/gr.113084.110

Li, H. (2011). A statistical framework for SNP calling, mutation discovery, association mapping and population genetical parameter estimation from sequencing data. Bioinformatics, 27(21), 2987–2993. https://doi.org/10.1093/bioinformatics/btr509

Li, H., & Durbin, R. (2010). Fast and accurate long-read alignment with Burrows-Wheeler transform. Bioinformatics (Oxford, England), 26(5), 589–595. https://doi.org/10.1093/bioinformatics/btp698

Liao, Y., Smyth, G. K., & Shi, W. (2014). featureCounts: An efficient general purpose program for assigning sequence reads to genomic features. Bioinformatics, 30(7), 923–930. https://doi.org/10.1093/bioinformatics/btt656

Libioulle, C., Louis, E., Hansoul, S., Sandor, C., Farnir, F., Franchimont, D., Vermeire, S., Dewit, O., de Vos, M., Dixon, A., Demarche, B., Gut, I., Heath, S., Foglio, M., Liang, L., Laukens, D., Mni, M., Zelenika, D., Van Gossum, A.,… Georges, M. (2007). Novel Crohn disease locus identified by genome-wide association maps to a gene desert on 5p13.1 and modulates expression of PTGER4. PLoS Genetics, 3(4), e58. https://doi.org/10.1371/journal.pgen.0030058

Love, M. I., Huber, W., & Anders, S. (2014). Moderated estimation of fold change and dispersion for RNA-seq data with DESeq2. Genome Biology, 15(12), 550. https://doi.org/10.1186/s13059-014-0550-8

Mackay, E. W., & Schulte-Merker, S. (2014). A statistical approach to mutation detection in zebrafish with next-generation sequencing. Journal of Applied Ichthyology, 30(4), 696–700.

Miesfeld, J. B., Gestri, G., Clark, B. S., Flinn, M. A., Poole, R. J., Bader, J. R., Besharse, J. C., Wilson, S. W., & Link, B. A. (2015). Yap and Taz regulate retinal pigment epithelial cell fate. Development, 142(17), 3021–3032. https://doi.org/10.1242/dev.119008

Minevich, G., Park, D. S., Blankenberg, D., Poole, R. J., & Hobert, O. (2012). CloudMap: A cloud-based pipeline for analysis of mutant genome sequences. Genetics, 192(4), 1249–1269. https://doi.org/10.1534/genetics.112.144204

Moffatt, M. F., Kabesch, M., Liang, L., Dixon, A. L., Strachan, D., Heath, S., Depner, M., von Berg, A., Bufe, A., Rietschel, E., Heinzmann, A., Simma, B., Frischer, T., Willis-Owen, S. A. G., Wong, K. C. C., Illig, T., Vogelberg, C., Weiland, S. K., von Mutius, E.,… Cookson, W. O. C. (2007). Genetic variants regulating ORMDL3 expression contribute to the risk of childhood asthma. Nature, 448(7152), 470–473. https://doi.org/10.1038/nature06014

Mullins, M. C., Hammerschmidt, M., Haffter, P., & Nüsslein-Volhard, C. (1994). Large-scale mutagenesis in the zebrafish: In search of genes controlling development in a vertebrate. Current Biology, 4(3), 189–202. https://doi.org/10.1016/S0960-9822(00)00048-8

Nicolae, D. L., Gamazon, E., Zhang, W., Duan, S., Dolan, M. E., & Cox, N. J. (2010). Trait-associated SNPs are more likely to be eQTLs: Annotation to enhance discovery from GWAS. PLoS Genetics, 6(4), e1000888.https://doi.org/10.1371/journal.pgen.1000888

Obholzer, N., Swinburne, I. A., Schwab, E., Nechiporuk, A. V., Nicolson, T., & Megason, S. G. (2012). Rapid positional cloning of zebrafish mutations by linkage and homozygosity mapping using whole-genome sequencing. Development, 139(22), 4280–4290. https://doi.org/10.1242/dev.083931

Poplin, R., Ruano-Rubio, V., DePristo, M. A., Fennell, T. J., Carneiro, M. O., Van der Auwera, G. A., Kling, D. E., Gauthier, L. D., Levy-Moonshine, A., Roazen, D., Shakir, K., Thibault, J., Chandran, S., Whelan, C., Lek, M., Gabriel, S., Daly, M. J., Neale, B., MacArthur, D. G., & Banks, E. (2017). Scaling accurate genetic variant discovery to tens of thousands of samples [Preprint]. Genomics. https://doi.org/10.1101/201178

R Core Team. (2019). R: A language and environment for statistical computing. R Foundation for Statistical Computing, Vienna, Austria. URL https://www.R-project.org/Stickney,

H.L., Schmutz, J. Woods, I. G., Holtzer, C. C., Dickson, M. C., Kelly, P. D., Myers, R. M., & Talbot, W. S. (2002). Rapid Mapping of Zebrafish Mutations With SNPs and Oligonucleotide Microarrays. Genome Research, 12(12), 1929–1934. https://doi.org/10.1101/gr.777302

Storey, J. D., Akey, J. M., & Kruglyak, L. (2005). Multiple locus linkage analysis of genomewide expression in yeast. PLoS Biology, 3(8), e267. https://doi.org/10.1371/journal.pbio.0030267

The 1000 Genomes Project Consortium. (2010). A map of human genome variation from population-scale sequencing. Nature, 467(7319), 1061–1073. https://doi.org/10.1038/nature09534

Thévenin, A., Ein-Dor, L., Ozery-Flato, M., & Shamir, R. (2014). Functional gene groups are concentrated within chromosomes, among chromosomes and in the nuclear space of the human genome. Nucleic Acids Research, 42(15), 9854–9861. https://doi.org/10.1093/nar/gku667

Tischler, G., & Leonard, S. (2014). biobambam: Tools for read pair collation based algorithms on BAM files. Source Code for Biology and Medicine, 9(1), 13. https://doi.org/10.1186/1751-0473-9-13

Wali, N., White, R. J., & Busch-Nentwich, E. M. (2021). RNA extraction from individual zebrafish embryos. Bio-Protocol. bio-protocol.org/prep873

White, R. J., Collins, J. E., Sealy, I. M., Wali, N., Dooley, C. M., Digby, Z., Stemple, D. L., Murphy, D. N., Billis, K., Hourlier, T., Füllgrabe, A., Davis, M. P., Enright, A. J., & Busch-Nentwich, E. M. (2017). A high-resolution mRNA expression time course of embryonic development in zebrafish. ELife, 6, e30860. https://doi.org/10.7554/eLife.30860

Zhang, J., Talbot, W. S., & Schier, A. F. (1998). Positional Cloning Identifies Zebrafish one-eyed pinhead as a Permissive EGF-Related Ligand Required during Gastrulation. Cell, 92(2), 241–251. https://doi.org/10.1016/S0092-8674(00)80918-6

